# FAPM: Functional Annotation of Proteins using Multi-Modal Models Beyond Structural Modeling

**DOI:** 10.1101/2024.05.07.593067

**Authors:** Wenkai Xiang, Zhaoping Xiong, Huan Chen, Jiacheng Xiong, Wei Zhang, Zunyun Fu, Mingyue Zheng, Bing Liu, Qian Shi

## Abstract

Assigning accurate property labels to proteins, like functional terms and catalytic activity, is challenging, especially for proteins without homologs and “tail labels” with few known examples. Unlike previous methods that mainly focused on protein sequence features, we use a pretrained large natural language model to understand the semantic meaning of protein labels. Specifically, we introduce FAPM, a contrastive multi-modal model that links natural language with protein sequence language. This model combines a pretrained protein sequence model with a pretrained large language model to generate labels, such as Gene Ontology (GO) functional terms and catalytic activity predictions, in natural language. Our results show that FAPM excels in understanding protein properties, outperforming models based solely on protein sequences or structures. It achieves state-of-the-art performance on public benchmarks and in-house experimentally annotated phage proteins, which often have few known homologs. Additionally, FAPM’s flexibility allows it to incorporate extra text prompts, like taxonomy information, enhancing both its predictive performance and explainability. This novel approach offers a promising alternative to current methods that rely on multiple sequence alignment for protein annotation. The online demo is at: https://huggingface.co/spaces/wenkai/FAPM_demo.

## Introduction

Proteins are essential components of cells and tissues, as they play key roles in various biological processes that sustain life. Understanding their functions is vital for decoding the molecular intricacies of biological systems and curing disease. Acquiring protein sequences from natural sources is generally straightforward, meanwhile, advancements in computational techniques, such as AlphaFold(*1*), have significantly improved the precision of predicting their three-dimensional structures(*2*). However, only fewer than 1% of the proteins, are accompanied by reviewed functional annotations(*3*) in global sequence databases such as UniProt(*4*). The majority that remains, with their functions still need to be experimentally characterized, are subject to high costs and logistical challenges. Furthermore, proteins demonstrate intricate variability and interactions as parts of their inherent complexity for their experimental validations. Therefore, predicting protein functions, while still challenging due to their complexity, is an important approach to elucidate their roles in pioneering studies.

Currently, protein functions can be described using the Gene Ontology (GO) (*5*), which is recognized as one of the most successful ontologies in biology. GO encompasses three sub-ontologies: molecular functions (MFO) characterizing the functions of individual proteins, biological processes (BPO) in which proteins participate, and cellular components (CCO) where proteins are active. The GO system facilitates a standardized approach to catalog the vast diversity of protein activities across different organisms. And the functions of these proteins are initially obtained from experimental evidences that published in the scientific literatures. These findings, once validated, are curated by database administrators into annotations within databases like UniProtKB/Swiss-Prot(*3*), ensuring that the information is accessible and standardized according to GO terms. Currently, the UniProtKB/Swiss-Prot database, known for its manually curated entries, encompasses GO annotations for over 560,000 proteins across thousands of organisms.

Various protein sequence-based methods have been proposed for predicating functions of proteins. As early in 1990, tools like BLAST(*6*) were developed, which allow researchers to compare an unknown protein sequence against a database of sequences with known annotations to find potential sequence similarities. The underlying assumption is that homologous sequences (those sharing a common evolutionary origin) will be of similar functions. Later, signature-based approaches(*7–9*) were introduced, which catalog domains and motifs within the proteins with similar functions based on scoring against large databases of statistical models for each sequence family with experimentally verified functions. These methods are now employed to update widely-used databases like InterPro(*10*). Despite their significant impact, these computational modeling approaches primarily rely on sequence similarity for prediction, leaving a considerable number of proteins unannotated.

Recent research has introduced a variety of deep neural network models to harvest multiple data simultaneously for functional predictions of protein, leveraging diverse sources of information, including sequence data, interactions and domain annotations. For example, DeepGOPlus(*11*), DeepGOZero(*12*) and DeepGraphGO(*13*) utilize domain features extracted from InterPro to train a classification model. In particular, DeepGraphGO uses graph convolutional layers(*14*) to capture neighboring information within the protein-protein interaction (PPI(*15*)) network extracted from STRING(*16*). Inspired by the success of convolutional neural networks(CNN)(*17*) in image, DeepGOCNN(*11*), DeepGOPlus and ProteInfer(*18*) utilize convolutional layers in CNN to extract features from protein sequences. Notably, ProteInfer inputs each amino acid after one-hot coding into the CNN network structure. Transformers(*19*) have achieved great success in many artificial intelligence fields(*20, 21*). Recently, it has also been discovered to have potential in protein modeling. ProteinBert(*22*) utilises protein sequences and semantic features of GO tags for pre-training. GoProFormer(*23*) combines sequence transformer and graph transformer to learn sequence features and GO network features for predicting GO tags. ESM-2 (Evolutionary Scale Modeling-2)(*24*) learns patterns and properties in protein sequences through pre-training on a large number of unlabeled protein sequences. These trained models are then applied to specific tasks of predicting protein structure and properties using fine-tuning and transfer learning, leading to improved performance in biomedical research, drug design, and other related fields(*25–27*).

Many large language models (LLMs) are capable of multitasking, including tasks related to proteins. For example, Galactica(*28*) is a LLM trained on scientific knowledge corpus, which can be applied to multi-modal tasks involving SMILES chemical formulas and protein sequences. And Mol-Instructions(*29*) combine construction methods of self-instruct, template-based conversion, and human-crafted task descriptions to cover a wider range of biomolecular tasks. The methods mentioned above primarily depend on sequence similarity or sequence feature modeling. However, the semantic information inherent in the functional description of proteins is rarely used, which is key to establish a comprehensive sequence-to-function link.

Here, we use a contrastive learning framework to jointly model protein sequences and functional texts in our model. To be specific, we consider protein sequences as a modality, and the natural language of functional annotations as another modality. We align the protein sequence modality to natural language modality using contrastive learning approach. This leads to our framework FAPM (**F**unctional **A**nnotation of **P**roteins using **M**ulti-modal models) shown in Figure 1, a contrastive learning method that generates functional descriptions for proteins by leveraging a pre-trained protein sequence model and large language model. FAPM efficiently harnesses the generative capabilities of language models to articulate and generate the functional-semantic information inherent in protein sequences. The notable features of our work are as follows:

1. Through the integration of multi-modal modeling, we effectively combine protein sequence modeling with language generation frameworks. The analysis in “the explainability of multi-modal representation” section shows the natural connection between the learned representations and their property descriptions, allowing for the direct generation of accurate annotations.
2. Comparative evaluations on the Swiss-Prot dataset and dozens of phage proteins demonstrate FAPM’s superior predictive performance over existing methods, underscoring the model’s effectiveness in accurately predicting protein functions.
3. Optional information, such as taxonomy, can be used as the additional input for the model to improve the generation quality, demonstrating a more flexible way to annotate proteins.

**Figure 1.**
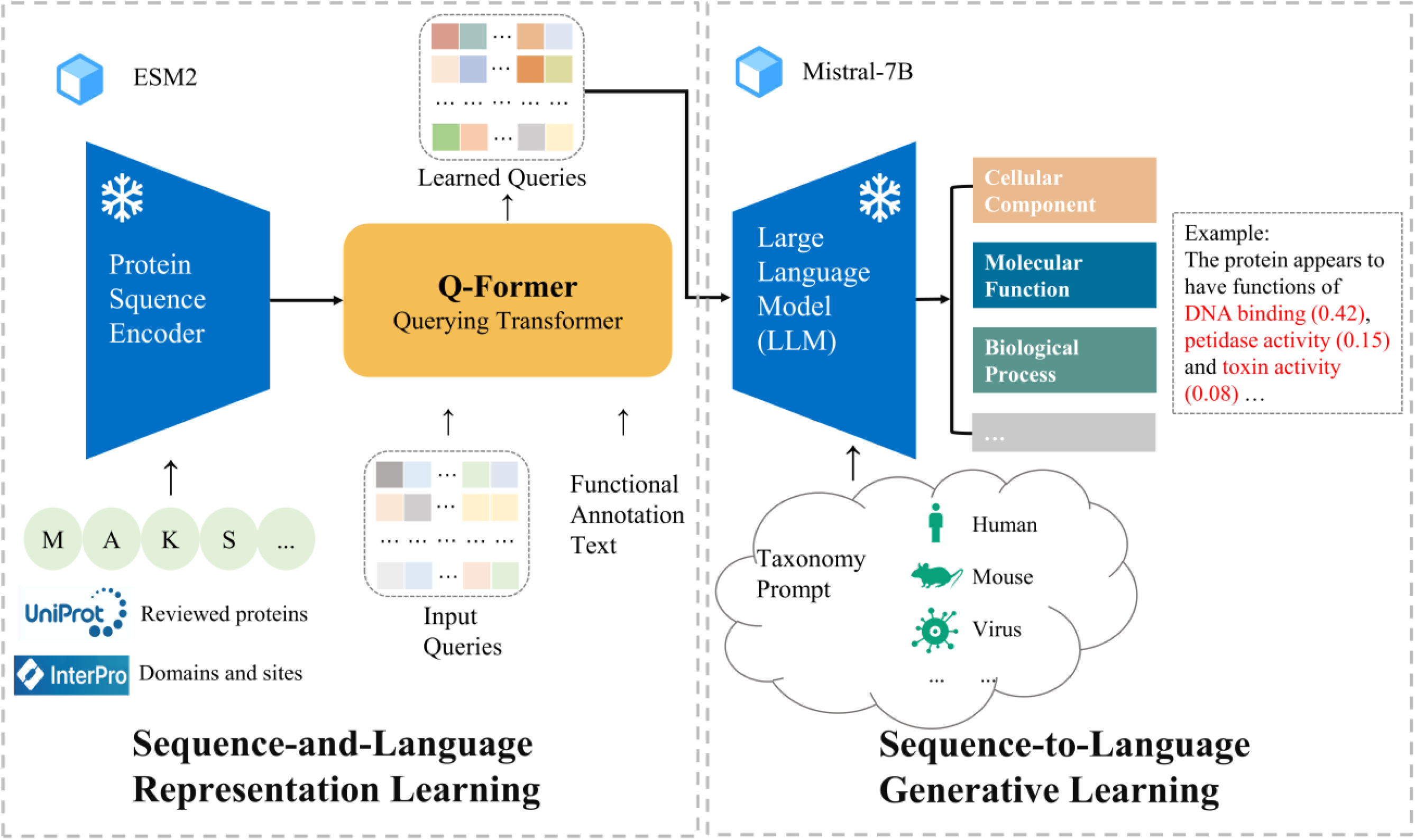
The overview of the framework of FAPM. In the Sequence-and-Language Representation Learning stage, FAPM use ESM2 to encode protein sequence and Q-Former to align them with function text (GO terms). In the Sequence-to-Language Generative Learning stage, Mistral-7B processes the learned queries from the prior stage, optionally incorporating prompts (taxonomy), to generate GO term predictions (MFO, BPO and CCO) with associated probabilities. The protein domain dataset from the InterPRO database and reviewed proteins from the UniProt database are utilized to enhance prediction performance.

## Results

We evaluate FAPM’s overall performance through two comparisons: (1) against other language models for multi-task prediction in the Instruction dataset (see “Datasets” section), and (2) with other popular methods, including InterPRO, DeepGOCNN, DeepGraphGO, DeepGOZero, DeepGO-SE, ProteInfer, ProtNLM, DeepFRI and AnnoPRO, on functional annotation tasks. These competing methods have been briefly described in the “Competing methods” section. Unfortunately, limitations in model open-source availability and training resources prevented us from reproducing some methods or making large-scale predictions, leading to the unavailability of test results on the Swiss-Prot test set. we conducted testing of these methods using a collection of bacteriophage and their bacterial host proteins. Since the experimental annotations for these bacteriophage/bacteria proteins have not been publicly updated, ensuring data integrity, a fair comparison could be conducted.

### Comparative Performance Against Other Language Models

Since the FAPM framework incorporates a language model, a straightforward idea is to compare FAPM with other language models to evaluate its performance in protein prediction tasks. We leverage the protein-oriented instruction dataset along with its train/test split to train our model from the beginning. Figure 2 (A) demonstrates the rouge-L metrics of eight methods across four tasks: protein function, functional description, catalytic activity and domain/motif prediction. The protein instance “UMP-CMP kinase” in Figure 2 (B) highlights that FAPM exhibits fewer errors in the generation, better alignment with the ground truth in functional descriptions, and successfully predicts functions (’phosphorylation’ and ‘nucleus’) that the second-best method (Mol-Instructions) failed to capture.

**Figure 2.**
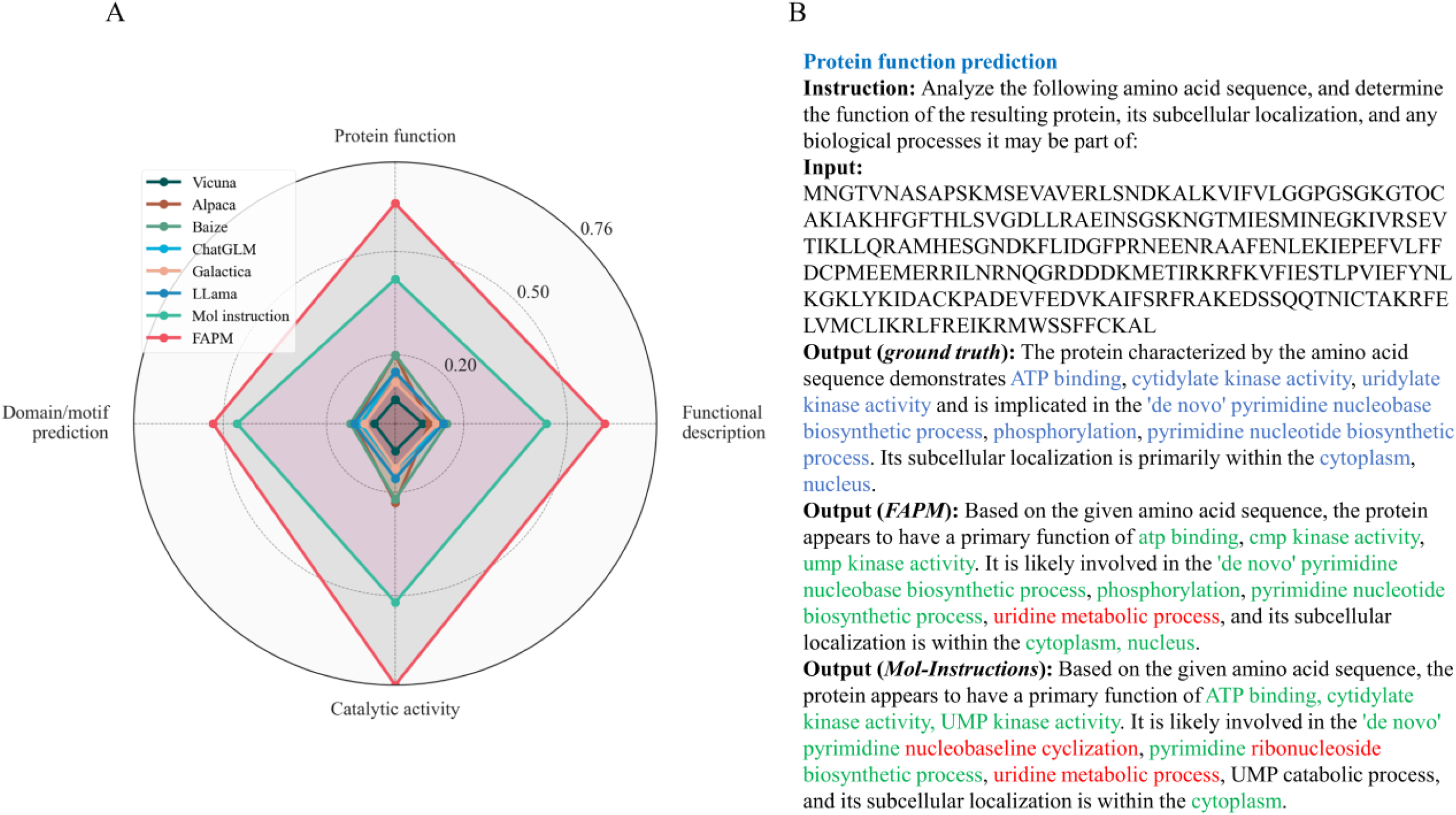
Performance against other language models. (**A**) The performance comparison on protein understanding tasks. (**B**) Examples of predicting protein function using instructions.

### Comparison with competing methods on the Swiss-Prot dataset

As shown in table 1, we made a comparison of FAPM with competing methods: Naive, DeepGOCNN, DeepGraphGO and DeepGOZero. It’s important to note that due to open-source or computing resource limitations, ProteInfer, DeepFRI and AnnoPRO cannot be retrained, while ProtNLM is primarily used for predicting protein names and does not easily compute evaluation metrics. As a result, we did not compare these methods in this section. According to table 1, we have five findings:

1. FAPM_ESM2-3B+prompt_ has the best performance of F_max_, WF_max_, AUPR and S_min_ in MFO, BPO and CCO. For example, it achieved the highest F_max_ of 0.586 in MFO, which was 5.2% improvement over DeepGO-SE (0.534).
2. FAPM_ESM2-3B+prompt_ achieves relatively small advantage across some metrics in the BPO domain, with 2.4% improvement over DeepGraphGO in AUPR. The explanation can be inferred from Figure 4(A): when the recall value is approximately between 0 and 0.3, the precision value of DeepGraphGO is significantly higher than that of other methods, leading to its higher AUPR value. This could be attributed to its effective utilization of PPI network information(*13*).
3. Compared to the Naive method, other models achieve a more significant advantage in the WF_max_ metric other than F_max_, as improved predictive performance for rare annotations leads to higher WF_max_ scores. For example, in CCO, the F_max_ of DeepGOZero (0.593) is close to Naive (0.565), however, the WF_max_ of DeepGOZero (0.474) is far more than Naive (0.351). As we can see, FAPM achieved both advantages in F_max_ and WF_max_, showing its proficiency in predicting both common and rare GO annotations.
4. The sequence-based deep learning method, DeepGOCNN, performed poorly across all three GO domains. This suggests that using a basic convolutional neural network to encode protein sequences may struggle to capture the most relevant information for protein function prediction.
5. Compared to FAPM_ESM2-650M_, FAPM_ESM2-650M+prompt_ performs better, especially in the bp and cc domain. We guess that the species information in the prompt is related to special functions. The case of homologues proteins with distinct functions in Figure 3 supports our suspicions. Inputting taxonomy (such as “Homo”) in the form of a prompt as exterior supplementary to the language side of FAPM, we monitored the changes of the model’s generated abilities. Encouragingly, we noticed improvements in the generated content, such as avoiding some basic errors and enhancing the quality of our predictions. For instance, considering a pair of homologous proteins, TRPV1_HUMAN and A0A452GR26_9SAUR, categorized as Homo and Phasianus (a type of turkey) in the taxonomic hierarchy. They exhibit a high degree of similarity, with a sequence similarity of 70.76% and a structural RMSD value down to 1.593. However, a crucial difference lies in their abilities to function as capsaicin receptors (TRPV1), which is a heat-activated cation channel that is modulated by inflammatory agents and contributes to acute and persistent pain(*30*). As we know, the seeds of Capsicum plants are dispersed predominantly by birds. In birds, the TRPV1 channel does not respond to capsaicin or related chemicals but mammalian TRPV1 is very sensitive to it, in contrast, mammalian TRPV1 channels are responsive to capsaicin due to their functional diversity and selective sensitivity, resulting in a spicy taste sensation. The function of TRPV1 is associated with “behavioral response to pain” (GO:0048266) in the GO system. With the species information, our model correctly annotated TRPV1_HUMAN with GO:0048266 while not for A0A452GR26_9SAUR, which is consistent with the aforementioned knowledge.

**Figure 3.**
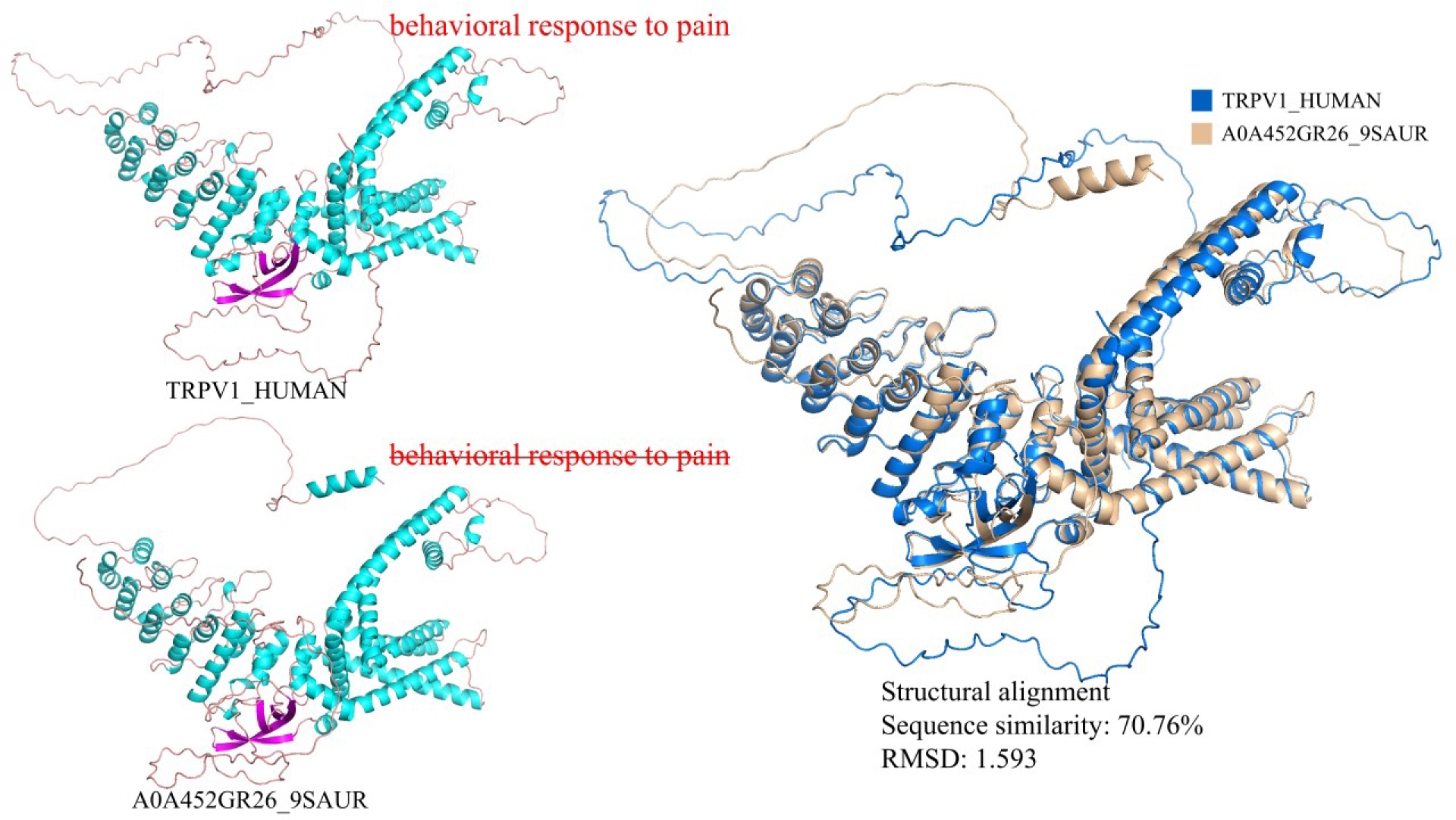
Case of functional prediction of homologous proteins from different species.

**Figure 4.**
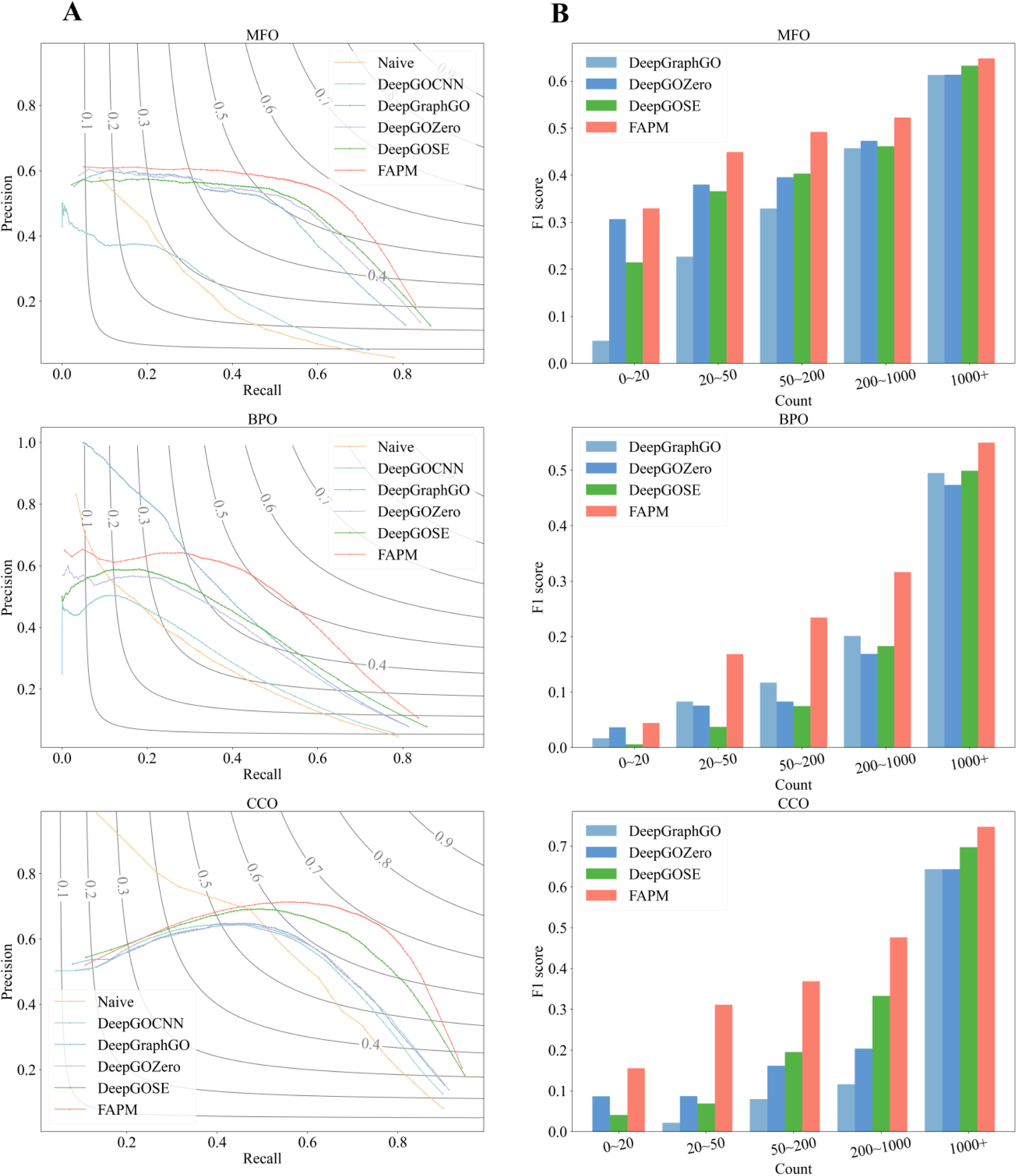
Performance against other models. (**A**) Recall-precision curve of various methods for Molecular Function, Biological Process and Cellular Component prediction on the test set of UniProtKB Swiss-Prot dataset. (**B**) The relationship between the frequency of Gene Ontology (GO) labels on the training set and the F1 scores of predictions on the test set of UniProtKB Swiss-Prot dataset.

**Table 1.**
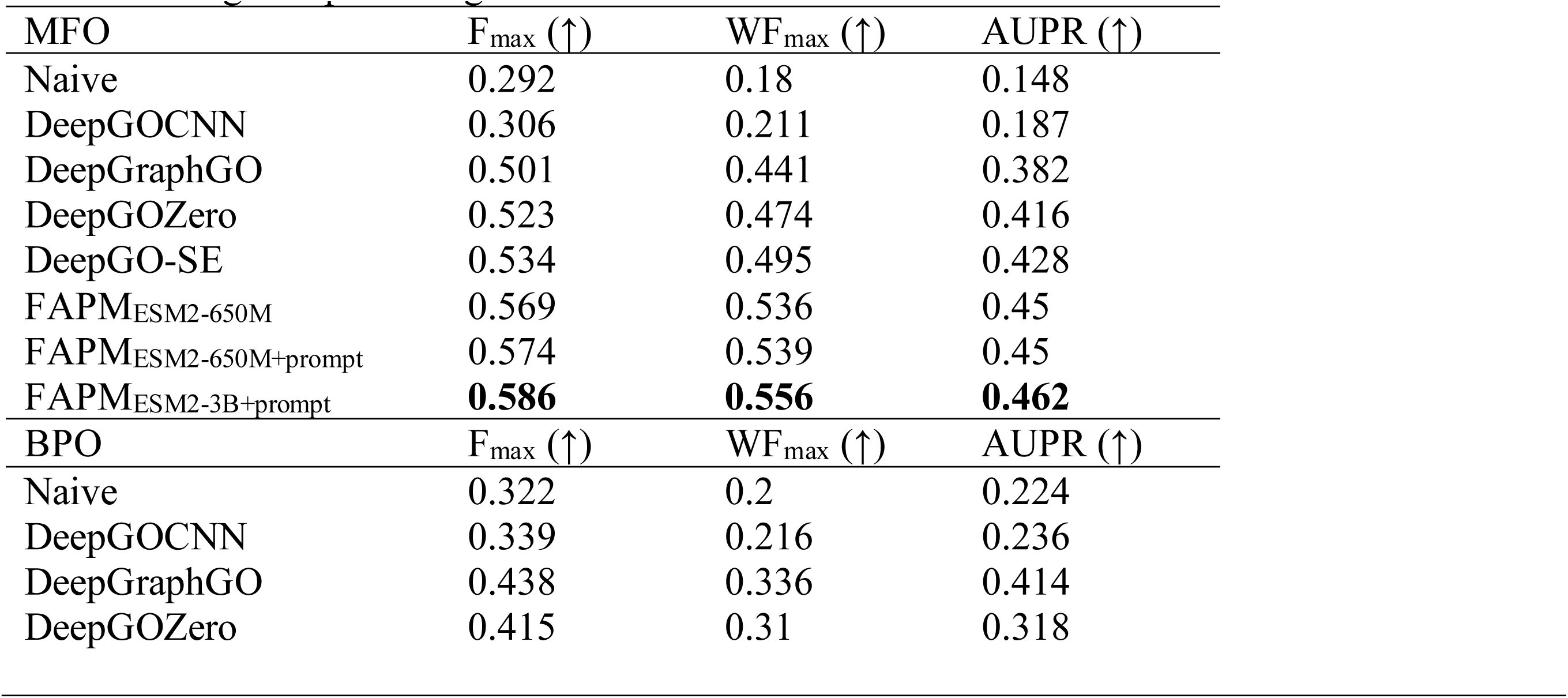
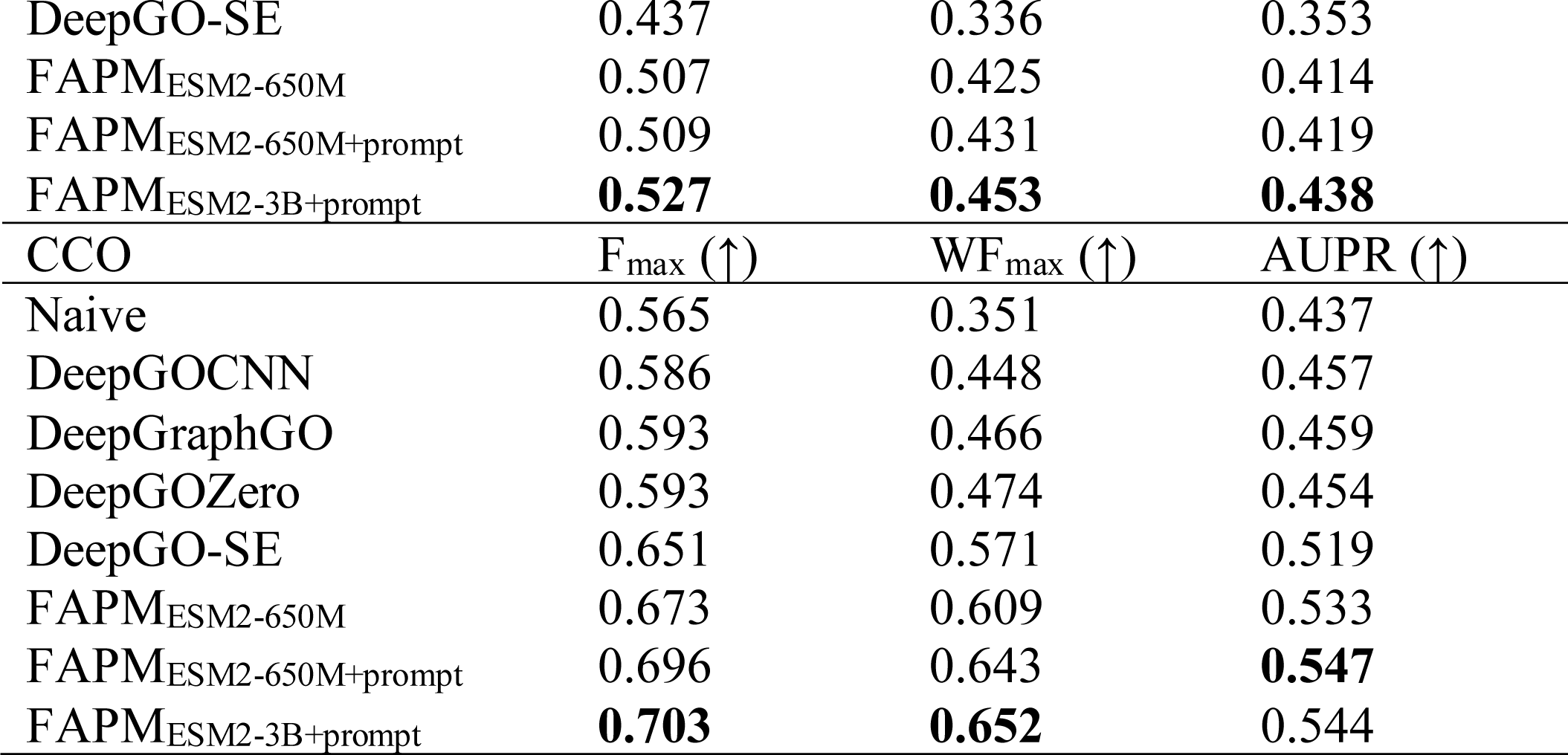
Prediction results for MFO, BPO and CCO on the test set of UniProtKB Swiss-Prot dataset.

| MFO | $F_{\max}$ (↑) | $WF_{\max}$ (↑) | AUPR (↑) |
| --- | --- | --- | --- |
| Naive | 0.292 | 0.18 | 0.148 |
| DeepGOCNN | 0.306 | 0.211 | 0.187 |
| DeepGraphGO | 0.501 | 0.441 | 0.382 |
| DeepGOZero | 0.523 | 0.474 | 0.416 |
| DeepGO-SE | 0.534 | 0.495 | 0.428 |
| FAPM<sub>ESM2-650M</sub> | 0.569 | 0.536 | 0.45 |
| FAPM<sub>ESM2-650M+prompt</sub> | 0.574 | 0.539 | 0.45 |
| FAPM<sub>ESM2-3B+prompt</sub> | **0.586** | **0.556** | **0.462** |
| BPO | $F_{\max}$ (↑) | $WF_{\max}$ (↑) | AUPR (↑) |
| Naive | 0.322 | 0.2 | 0.224 |
| DeepGOCNN | 0.339 | 0.216 | 0.236 |
| DeepGraphGO | 0.438 | 0.336 | 0.414 |
| DeepGOZero | 0.415 | 0.31 | 0.318 |
| DeepGO-SE | 0.437 | 0.336 | 0.353 |
| FAPM<sub>ESM2-650M</sub> | 0.507 | 0.425 | 0.414 |
| FAPM<sub>ESM2-650M+prompt</sub> | 0.509 | 0.431 | 0.419 |
| FAPM<sub>ESM2-3B+prompt</sub> | **0.527** | **0.453** | **0.438** |
| CCO | $F_{\max} (\uparrow)$ | $WF_{\max} (\uparrow)$ | AUPR (↑) |
| Naive | 0.565 | 0.351 | 0.437 |
| DeepGOCNN | 0.586 | 0.448 | 0.457 |
| DeepGraphGO | 0.593 | 0.466 | 0.459 |
| DeepGOZero | 0.593 | 0.474 | 0.454 |
| DeepGO-SE | 0.651 | 0.571 | 0.519 |
| FAPM<sub>ESM2-650M</sub> | 0.673 | 0.609 | 0.533 |
| FAPM<sub>ESM2-650M+prompt</sub> | 0.696 | 0.643 | **0.547** |
| FAPM<sub>ESM2-3B+prompt</sub> | **0.703** | **0.652** | 0.544 |

The precision-recall curves of six methods on the test set of UniProtKB-SwissProt dataset are plotted in Figure 4(A). It is worth noting that DeepGOCNN did not outperform the Naive method by much, while other deep learning methods significantly surpassed the Naive method, especially in the MFO domain. Interestingly, in the BPO domain, if one aims for high precision in prediction results (at the cost of low recall), then DeepGraphGO is preferable. However, FAPM tends to perform better in general. In the CCO domain, the precision-recall curve of the Naive method crosses with that of other methods, indicating its prediction advantage in some cases (when recall is around or below 0.5).

Figure 4(B) illustrates the relationship between the frequency of GO term occurrences and the predictive performance of the models. We counted the occurrences of each GO term on the training set and grouped them into five categories: 0-20, 20-50, 50-200, 200-1000, and 1000+. The average F1 score for the predicted GO terms in each group was then calculated to compare the performance of four models: DeepGraphGO, DeepGOZero, DeepGO-SE and FAPM. When predicting high-frequency GO terms (1000+), the three methods show similar performance, with FAPM slightly ahead. However, for GO terms occurring less than 1000 times, FAPM demonstrates a more significant advantage, especially in BPO and CCO categories, highlighting FAPM’s strength in predicting rare GO terms.

### The explainability of multi-modal representation

FAPM achieves better quality protein representations in terms of properties compared to pretrained protein language model such as ESM2. Figure 5 visualizes the 2D projections of proteins with specific functions from the test set of the Swiss-Prot dataset using UMAP(*31*). The figure consists of three parts (A, B, and C), each comparing FAPM to ESM2. Part A focuses on most common functions (proteins with multi functions removed), while parts B and C show proteins annotated with similar and contrasting functions, respectively. In all three scenarios, the representations derived from ESM2 exhibit weak clustering, whereas FAPM yields significantly distinct and intriguing results. Specifically, part B (right) presents two closely positioned clusters, and part C (right) presents two relatively wider separated clusters, demonstrating that our model effectively captures the functional relationships of protein sequences. Overall, the representations learned by FAPM not only outperform those of ESM2 but also align with intuitive expectations about cluster distances, suggesting that our model successfully incorporates semantic information from property descriptions.

**Figure 5.**
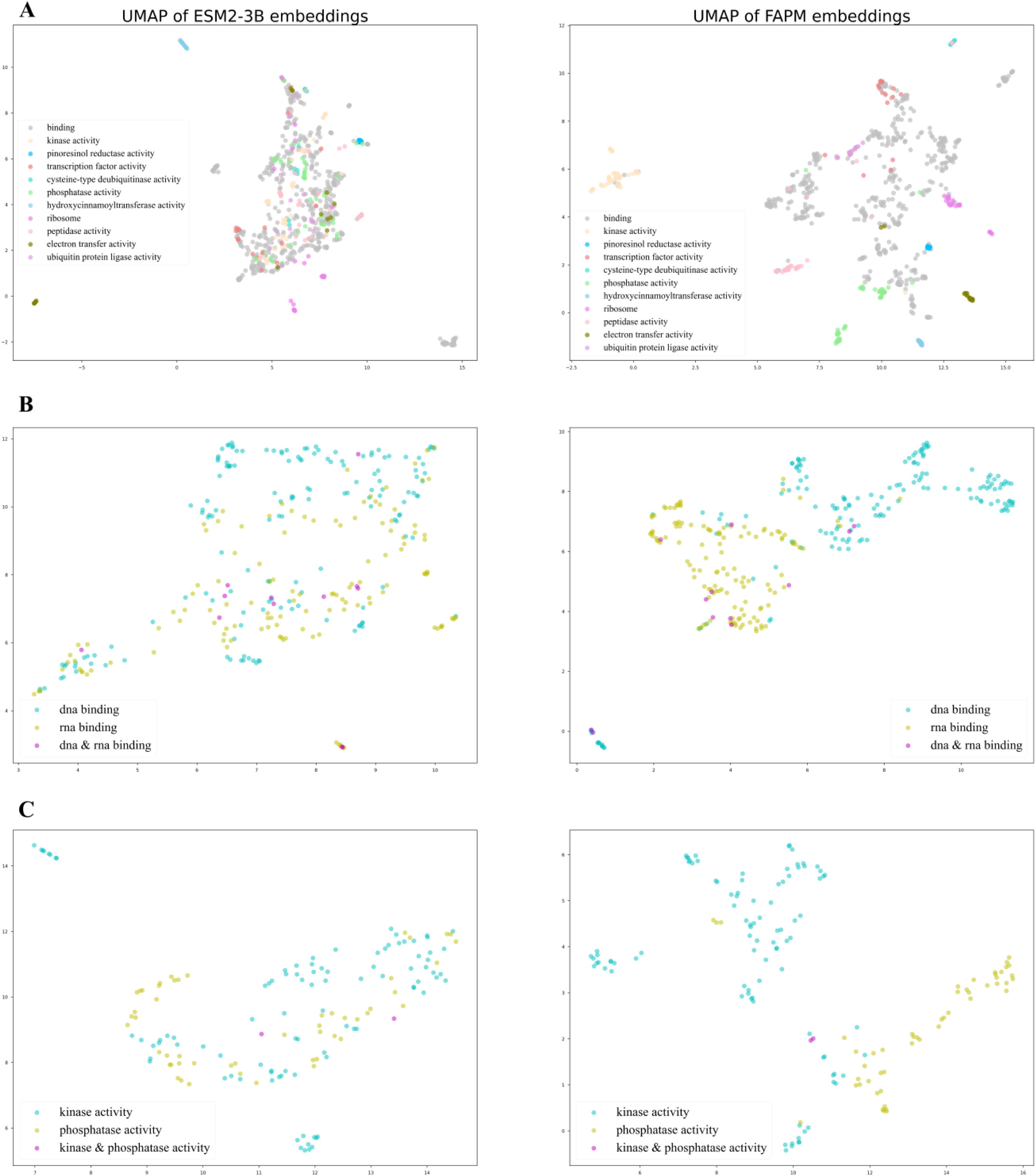
The comparisons of protein representations with specific GO functions between ESM2 and FAPM. All samples are from test set of Swiss-Prot dataset. (A) proteins annotated with most common functions. (B) proteins annotated with DNA/RNA binding (similar functions). (C) proteins annotated with kinase/phosphatase activity (contrast functions).

### Bacteriophage protein prediction cases

Predicting the functions of proteins without close homologs is challenging for both homology search and AI-based methods. To fully exploit the potential of our model, we chose bacteria and bacteriophage (phage) proteins to experimentally validate our predictions. Bacteria are the leading cause of pathogenic diseases and commensal microorganisms of human. Our understandings on bacteria are largely based on model bacteria, yet Escherichia coli (E. coli)(*32*) and Bacillus subtilis (B. subtilis)(*33*), representing Gram-negatives and Gram-positives, still requires vigorous further investigation as more than half of their genes still need functional assignment. Phages have an estimated population of 10^31^(*^34^*), more than all other lives combined on earth. The enormous population and variety of phage make their gene products largest reservoir of proteins with unknown functions, which are impossible to be annotated manually based on current experimental skill sets. We selected a wide range of bacteria and phage proteins, including those from E. coli, B. subtilis, phages for common bacteria, and rare RNA phage. Out of the 29 proteins (24 from phage and 5 from bacteria) selected, 3 recently assigned, and 5 are experimental validated but not described in the database uniport (Supplementary Table S1).

we evaluated FAPM against other six methods in predicting experimental GO annotations of bacteriophage proteins that mostly have no homology information available. Our analysis steps are as follows:

1. For these proteins, we conducted a search in the InterPRO database to obtain homology information, and also assessed their similarity with sequences in Swiss-Prot. This analysis aids in understanding the predictability of the sequence. For example, in InterPRO we can search that Gp57B has been annotated IPR009097 (Cyclic phosphodiesterase), which is defined as β-barrel domain consisting of a duplication of a β/α/β/α/β motif found in cyclic phosphodiesterase (CNPase)(*35–37*). And this definition can be considered as GO:0016787 (hydrolase activity). For similarity-based algorithms and AI algorithms, proteins with identifiable homology information, like Gp57B, are relatively easier to predict. However, there are many other proteins lacking homology information, such as Gp27, present a greater challenge for function prediction.
2. We employ various methods to predict protein functions, which can be classified into three categories. The first category consists of a single method that utilizes the homology search results from InterPRO for prediction, serving as our baseline method. The second category includes DeepGOZero, which are based on sequence annotation. We excluded DeepGOCNN due to its poor performance and DeepGraphGO because of the lack of protein interaction data. The third category focuses on purely sequence or structure-based methods, such as ProteInfer, ProtNLM, DeepFRI and AnnoPRO, all of which are trained on large scale datasets.
3. It’s important to note that, DeepGOZero, DeepGO-SE, ProtNLM, and AnnoPRO provide probabilities for nearly all GO labels, prompting us to select the top 10 predictions based on their scores. ProteInfer and DeepFRI only output the most likely predictions, so we did not further filter their predictions, which could be either more or fewer than 10.

We tabulated the occurrences of experimental annotations on the bacteriophage proteins, with “DNA binding” being the most frequent annotation, appearing 11 times. “RNA binding” appeared twice, while “Nuclease activity,” “Catalase activity,” and “Peptidase inhibitor activity” each appeared once. The recall, precision, and f1 score for these annotations across various methods are computed and presented in Table 2. However, due to the lack of predictive results of some methods for some GO labels, part of results is therefore not given.

**Table 2.**
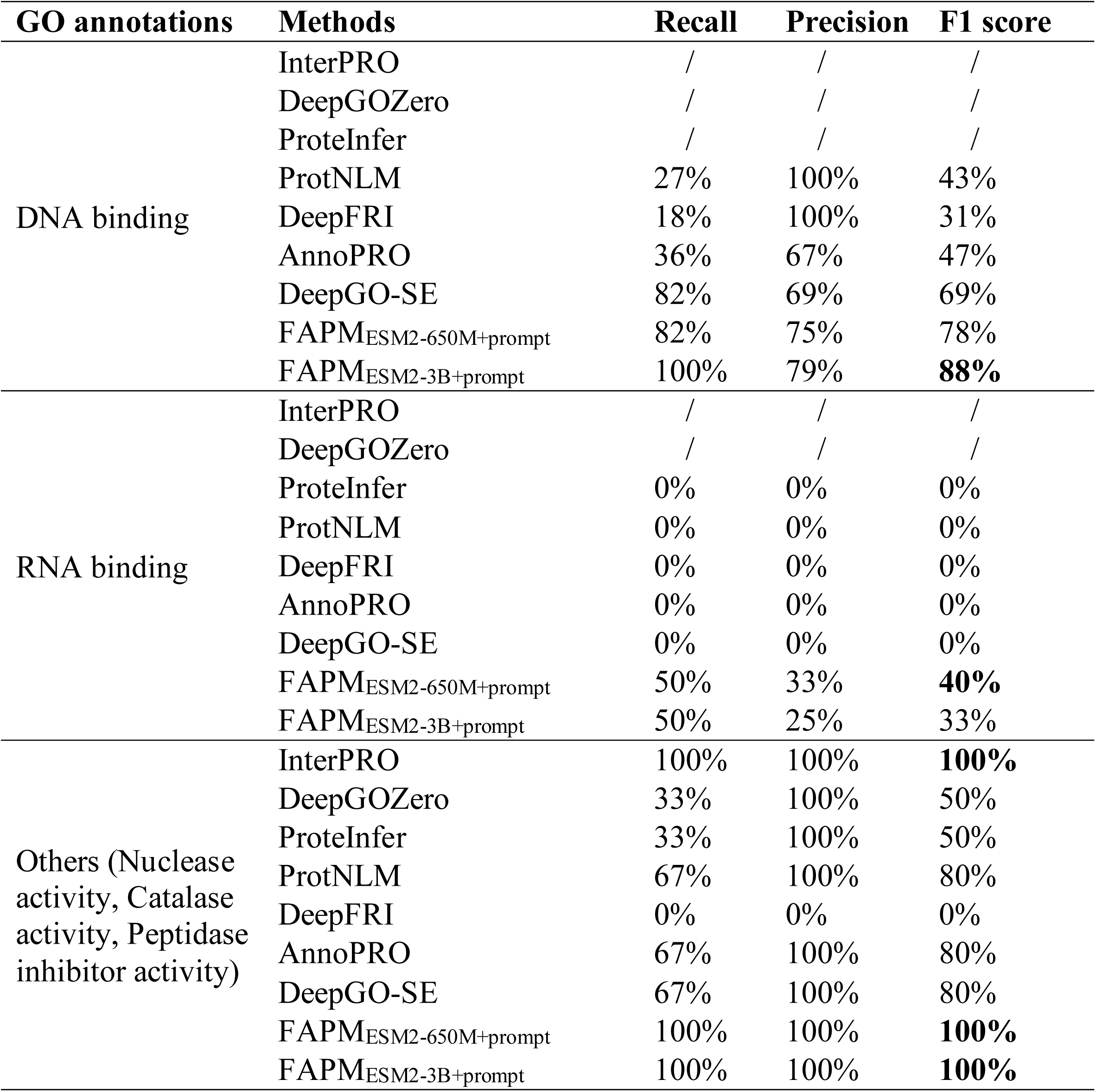
Recall, Precision and F1 Score of different methods for predicting GO terms of different bacteriophage proteins.

| GO annotations | Methods | Recall | Precision | F1 score |
| --- | --- | --- | --- | --- |
| DNA binding | InterPRO | / | / | / |
|  | DeepGOZero | / | / | / |
|  | ProteInfer | / | / | / |
|  | ProtNLM | 27% | 100% | 43% |
|  | DeepFRI | 18% | 100% | 31% |
|  | AnnoPRO | 36% | 67% | 47% |
|  | DeepGO-SE | 82% | 69% | 69% |
|  | FAPM<sub>ESM2-650M+prompt</sub> | 82% | 75% | 78% |
|  | FAPM<sub>ESM2-3B+prompt</sub> | 100% | 79% | **88%** |
| RNA binding | InterPRO | / | / | / |
|  | DeepGOZero | / | / | / |
|  | ProteInfer | 0% | 0% | 0% |
|  | ProtNLM | 0% | 0% | 0% |
|  | DeepFRI | 0% | 0% | 0% |
|  | AnnoPRO | 0% | 0% | 0% |
|  | DeepGO-SE | 0% | 0% | 0% |
|  | FAPM<sub>ESM2-650M+prompt</sub> | 50% | 33% | **40%** |
|  | FAPM<sub>ESM2-3B+prompt</sub> | 50% | 25% | 33% |
| Others (Nuclease activity, Catalase activity, Peptidase inhibitor activity) | InterPRO | 100% | 100% | **100%** |
|  | DeepGOZero | 33% | 100% | 50% |
|  | ProteInfer | 33% | 100% | 50% |
|  | ProtNLM | 67% | 100% | 80% |
|  | DeepFRI | 0% | 0% | 0% |
|  | AnnoPRO | 67% | 100% | 80% |
|  | DeepGO-SE | 67% | 100% | 80% |
|  | FAPM<sub>ESM2-650M+prompt</sub> | 100% | 100% | **100%** |
|  | FAPM<sub>ESM2-3B+prompt</sub> | 100% | 100% | **100%** |

For prediction of DNA binding, FAPM_ESM2-3B+prompt_ achieved F1 score of 88%, followed by DeepGO-SE with 69%. Notice that although both ProtNLM and DeepFRI have 100% precision, their recall values are too low, resulting in lower F1 scores. In the case of RNA binding, FAPM gave a successful prediction for Gp49, and none of the other methods gave a correct prediction for this annotation. The remaining 3 annotations (“Nuclease activity”, “Catalase activity” and “Peptidase inhibitor activity”) were successfully predicted by FAPM, whereas ProtNLM, AnnoPRO and DeepGO-SE made accurate predictions for two of them.

The test results from these cases demonstrate that FAPM exhibits consistent and outstanding performance in annotation prediction, even in some challenging cases.

## Discussion

FAPM utilizes a protein pre-training model, ESM2, to extract sequence information. And by aligning the sequence information with the natural language information, FAPM can use language model Mistral-7b to generate functional text. Therefore, it is a sequence-based protein function prediction model based on contrastive learning. In general, FAPM has three advancements in field:

1. Protein sequence is the only required input, and the output is easy to interpret due to the utilization of a language model.
2. Millions of protein domain data and hundreds of thousands of manually labeled data were used for model training. Our model demonstrates superior performance when compared to the alternative methods. While predicting the unannotated bacteriophage/bacteria proteins, our model exhibits better accuracy and generalization capabilities in predicting non-homologous proteins.
3. Users can interact conveniently with the model through prompt. In this study, we take taxonomy information as a kind of prompt, which significantly enhances the coherence of the output. Theoretically, more diverse prompts could be introduced to meet the demands of complex real-world scenarios.

For comparison and testing purposes, we have temporarily restricted the text generated by the model to the GO labelling system, which constrained the model’s flexibilities. However, FAPM holds significant potential for further development, with appropriate improvements, it may make richer predictions, including protein modification and generation. Moreover, enhancing the overall performance can be conveniently achieved by utilizing more powerful protein models and language models.

## Materials and Methods

FAPM utilizes a Querying Transformer (Q-Former) to address the modality gap in frozen pre-trained unimodal models. Q-Former are pre-trained in two stages: (1) protein-language representation learning stage with a frozen protein encoder and (2) protein-to-language generative learning stage with a frozen Large Language Model (LLM).

### Methods

#### Model Architecture overview

Figure 1 shows a schematic procedure of FAPM, two frozen pre-train models connected by Q-Former. Specially, we use ESM2 for protein sequence encoding and Mistral-7B(*38*) for text generation due to their outstanding performance. Q-Former, initialized with pre-trained weights of BERTbase(*39*), comprises transformer modules for extracting features from the frozen protein model and encoding/decoding the text. The total learnable parameters in Q-Former amount to approximately 188 million.

#### Protein Representation learning

In the representation learning stage, a raw amino acid sequence is first passed through ESM2 (3B), producing 2560-dimensional protein embeddings for each amino acid. Simultaneously, a set number of learnable query embeddings are generated as input to the Q-former along with the sequence embeddings. The queries will interact with frozen protein features through cross-attention structure and interact with input protein function text through self-attention structure. By default, the number of queries is set to 32, its size (32 × 768) is significantly smaller than the size of protein embeddings (sequence length × 2560) generated by ESM2 (3B). This approach enables the model to learn more compact representations while decreasing the total number of parameters.

#### Protein function generation learning

In the generation stage, we use LLM to generate functional labels such as GO terms. The GO file is of version 1.4 (http://purl.obolibrary.org/obo/go/go-basic.obo), ensuring that the graph is acyclic and allowing for the propagation of annotations up the graph. Specially, LLM processes learned queries from stage 1, along with an additional taxonomy prompt, and then generates text based on this information for molecular function, biological process, and cellular component. Note that the learned queries are preceded by a full connected network, and the taxonomy prompt can also be null.

In terms of the LLM option, instead of OPT-2.7B used in BLIP2, we choose Mistral-7B, a widely used open-source large language model developed by Mistral AI, as a core component of FAPM, due to its superior performance and better development ecosystem. Although Mistral-7B has over twice the number of parameters than OPT-2.7B, the frozen nature of these parameters does not impose a significant burden on the training and inference of the model.

We have included an additional input prompt, not primarily intended to enhance the model’s prediction accuracy. For instance, certain prompt categories (e.g., taxonomy) may not substantially improve model performance. Our goal is to introduce interactivity and flexibility to the model. In practical prediction scenarios, proteins are often accompanied by supplementary information, which we aim for the model to engage with and comprehend through prompts, resulting in more precise content. For instance, the inclusion of basic taxonomy information prompts can lead to more reasonable output content and reduce the likelihood of conflicting information.

#### Pre-Training Objectives

Inspired by BLIP2, we jointly optimize three pre-training objectives that share the same input format and model parameters.

**Protein-Text Contrastive Learning (PTC)** aims to align the feature space of the protein encoder and the text decoder, encouraging parallel protein-text pairs have higher similarity scores. This objective has been demonstrated to be effective in ALBEF(*40*) in image-text learning task.

**Language Modeling Loss (LM)** assess the efficacy of the language model to generate function text given a protein sequence. This metric quantifies the model’s proficiency in forecasting the likelihood of subsequent tokens in a sequence, based on the discrepancy between the predicted probabilities and the actual token occurrences.

**Protein-Text Matching (PTM)** predicts whether a pair of protein and text is positive (matched) or negative (not matched). We follow the negative sampling strategy(*40*), where negative pairs with higher contrastive similarity in a batch have a higher chance to be sampled. It aims to learn protein-text multimodal representation for protein-text pair that share similar semantics but differ in fine-grained details.

### Datasets

#### UniProtKB/Swiss-Prot Dataset

We downloaded the Swiss-Prot version 2023_04(*4*), released on October 25, 2023, from the FTP site (https://ftp.uniprot.org/pub/databases/uniprot/previous_releases/), which is a comprehensive resource of protein sequence and annotation information. We filtered sequences shorter than 10 or longer than 1024, then we divided these protein sequences into two parts, the portion automatically annotated was utilized as pre-training samples along with the domain-scale sequences, while the remaining part was used for fine-tuning. Experimental functional annotated proteins can be filtered with evidence codes EXP, IDA, IPI, IMP, IGI, IEP, TAS, IC, HTP, HDA, HMP, HGI and HEP, which contains 71239 reviewed and manually annotated proteins. To fine-tune our model on this dataset and guarantee effective generalization to new proteins, we partitioned the proteins into training, validation, and testing sets. We employed DIAMOND(*41*) to compute sequence similarity, grouping the proteins based on their similarity before executing a random split. Proteins with sequence identity exceeding 50% were grouped together, with 90% of the groups allocated for training, 5% for validation, and 5% for testing. Table 3 shows the number of samples under the sub-ontologies of GO in the pre-training and fine-tuning phase, as well as the number of GO terms.

**Table 3.**
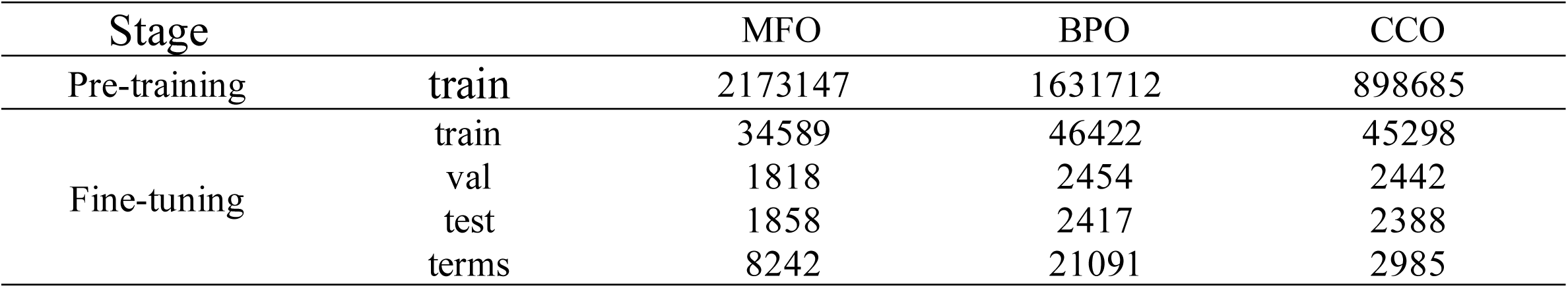
Summary of pre-training and fine-tuning dataset.

To improve the model’s ability to learn domain-scale information from sequence, we pre-train the model using domain data generated by InterPro. We utilized data version 95.0 (2023-07), available for download from the InterPro FTP site (https://ftp.ebi.ac.uk/pub/databases/interpro/). However, as these domain sequences are annotated with domain number (such as IPR000001), we utilize the undergoing mapping provided by InterPro to assign GO terms to these sequences.

#### Instruction dataset

We use the protein-oriented instructions introduced by Mol-Instructions (https://huggingface.co/datasets/zjunlp/Mol-Instructions), which covers 505K instructions spanning five categories of tasks. These tasks aim to predict the structure, function, and activity of proteins, and facilitate protein design based on textual directives. In this study, we exclude protein design task due to model limitation. Performance evaluation is conducted using the ROUGE-L(*42*) metric on all test samples to measure the quality of the model’s output against reference answers.

#### Competing methods

A variety of compared methods are summarized in Table 4, from simplest naïve methods to newest state-of-the-art method DeepGO-SE(*43*). AI-based methods mainly rely on protein features or protein sequences as inputs, supplemented with information like PPI or semantic sometimes.

**Table 4.** Summary of comparable methods.

| Methods | Input |
| --- | --- |
| Naive | Statistic information |
| InterPRO | Homology information |
| DeepGraphGO | Protein features and PPI information |
| DeepGOZero | Protein features and semantic information |
| DeepGO-SE | Protein features and semantic information |
| DeepGOCNN | Sequence |
| ProteInfer | Sequence |
| ProtNLM | Sequence |
| AnnoPRO | Sequence |
| DeepFRI | Sequence and structure |
| Mol-Instructions | Sequence |

### Naive baseline

In a straightforward scenario, we estimate the GO probability of each protein based on the frequency of GO occurrences in the data. This basic approach offers advantages in predicting high-frequency GO tags due to the imbalanced nature of GO category annotations and can serve as a baseline called “naive” method(*44*). The prediction scores for GO class 𝑓 in protein 𝑝 are computed as:

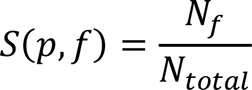

Where 𝑁_𝑓_ is the number of training proteins annotated by GO class 𝑓 and 𝑁_𝑡𝑜𝑡𝑎𝑙_ is a total number of training proteins.

### Other methods

- InterPRO is a resource that provides functional analysis of protein sequences by classifying them into families and predicting the presence of domains and important sites through collaboration with multiple member databases (referred to as member databases). Additionally, InterPro provides detailed functional annotations as well as adding relevant GO terms that enable automatic annotation of millions of GO terms across the protein sequence databases.
- DeepGOZero integrates protein function prediction with ELEmbeddings(*45*), a model-theoretic approach for embedding ontologies into a distributed space. It uses InterPro domain annotations as input and employs two layers of MLPBlock to generate embeddings, then the model learns the embedding space for GO classes using ELEmbeddings’ loss functions.
- DeepGraphGO is a multi-species graph neural network-based method for protein function prediction, designed to leverage sequence features (InterPRO domain annotations) and high-order protein network information in an end-to-end manner.
- ProteInfer predicts protein function directly from unaligned amino acid sequences using deep convolutional neural networks. This method could predict Enzyme Commission (EC) numbers and Gene Ontology (GO) terms, while also placing amino acid sequences into a functional space to assist in downstream analysis and interpretation.
- ProtNLM aimed to develop a model that connects amino acid sequences with natural language descriptions of their functional properties. This model was trained to predict free text captions describing these properties. Expert curators evaluated model predictions, leading to performance enhancements. The refined model was then used to predict protein names for around 49 million unknown proteins, now included in the UniProt database.
- DeepFRI(*46*) is a Graph Convolutional Network that predicts protein functions by utilizing sequence features from a protein language model and structures. DeepFRI uses LSTM(*47*) language model to extract residue-level features of PDB(*48*) sequence and GCN(*49*) with three graph convolutional layers to learn complex structure–function relationships.
- AnnoPRO(*50*) incorporates sequence-based multi-scale protein representation, dual-path protein encoding through pre-training, and function annotation using long short-term memory-based decoding. It aims to address the challenge of long-tail problem(*51*).
- DeepGO-SE predicts GO functions from protein sequences. It integrates a pre-trained protein language model with a neuro-symbolic model that leverages GO axioms to perform protein function prediction through approximate semantic entailment.

### Experimental settings

We train and evaluate models for each of the sub-ontologies of GO separately. Initially, our methodology employs millions of domain level sequences and several hundred thousand automatically annotated full-chain level sequences in pre-training phase. This phase encompasses a two-stage training process: a 5-epoch protein-text alignment training, followed by a 5-epoch training focused on generating protein functional text outputs. Subsequently, model selection is guided by the validation set’s performance metrics. Building upon the pre-trained model, we further refine its performance through a 5-epoch fine-tuning process, utilizing manually annotated data and adopting a lower learning rate of 1e-5.

### Evaluation metrics

To summarize model performance as a single scalar, we compute the F_max_, AUPR and WF_max_. F_max_, the maximum F1 score (the geometric mean of precision and recall) across all thresholds, is protein-centric, which has been used in CAFA as the main evaluation metric(*52*). AUPR, the area under the precision-recall curve, is pair-centric and widely used for performance evaluation of multi-label classification including automated function prediction(*11, 53*). WF_max_ is weighted version of F_max_, and the weight of GO annotation is calculated as information accretion (IA)(*54*).

## Author contributions

Conceptualization, Z.X., M.Z. and Q.S

Methodology, Z.X., B.L., M.Z., Q.S., W.X., X.H., J.X., Z.F. and W.Z.

Software, W.X. and Z.X. Investigation, W.X., Z.X. and Q.S.

Writing – Original Draft, W.X.

Writing – Review & Editing, Z.X., W.X., B.L., H.C. and Q.S.

Funding Acquisition, M.Z. and Q.S.; Resources, B.L. and H.C.

Supervision, Z.X., B.L., M.Z. and Q.S.

## Data and materials availability

Code:

https://github.com/xiangwenkai/FAPM

Data:

https://ftp.uniprot.org/pub/databases/uniprot/previous_releases/release-2023_04/knowledgebase/ https://huggingface.co/datasets/zjunlp/Mol-Instructions https://ftp.ebi.ac.uk/pub/databases/interpro/

Demo:

https://huggingface.co/spaces/wenkai/FAPM_demo

## Supplementary Materials

**Table S1.** The experimentally verified bacteriophage and bacteria proteins that are unannotated in UniProt. Proteins are bacteriophage protein unless listed otherwise.

| Name | location | Functional labels |
| --- | --- | --- |
| Gp41 | cytosol | DNA binding; protein-protein interaction |
| Gp44 | cytosol | DNA binding |
| Gp45 | cytosol | DNA binding |
| Gp46 | cytosol | Protein-protein interaction |
| Gp49 | cytosol | RNA binding |
| Gp60 | cytosol | Protein-protein interaction |
| Gp33 | cytosol | Protein-protein interaction |
| Gp27 | cytosol | DNA binding |
| Gp35.1 | cytosol | Toxin activity |
| RpbA | cytosol | Protein-protein interaction |
| Exod | cytosol | Dual nuclease activity |
| Gp57B | cytosol | Catalase activity |
| Pin | cytosol | Peptidase inhibitor activity; Protein-protein interaction |
| Mrh | cytosol | DNA binding; Protein-protein interaction |
| Cef | cytosol | RNA binding; Protein-protein interaction |
| MsyB | cytosol | Protein-protein interaction, E. coli |
| YciZ | cytosol | Signal transduction; Protein-protein interaction, B. subtilis |
| YlaB | inner membrane | Signal transduction; Protein-protein interaction, B. subtilis |
| Ysdb | inner membrane | Signal transduction; Protein-protein interaction, B. subtilis |
| RsiW | transmembrane | Signal transduction; Protein-protein interaction, B. subtilis |
| Gp2 | cytosol | Protein-protein interaction |
| Gp6 | cytosol | DNA binding |
| Gp8 | cytosol | DNA binding |
| Gp12 | cytosol | DNA binding |
| ss1 |  | DNA binding |
| sm2 | cytosol | Protein-protein interaction |
| sl1 | cytosol | Protein-protein interaction |
| 75 | cytosol | DNA binding |
| 235 | cytosol | DNA binding |

## Reference

1. J. Jumper et al., Highly accurate protein structure prediction with AlphaFold. Nature 596, 583–589 (2021).

2. M. Varadi et al., AlphaFold Protein Structure Database: massively expanding the structural coverage of protein-sequence space with high-accuracy models. Nucleic acids research 50, D439–D444 (2022).

3. B. Boeckmann et al., The SWISS-PROT protein knowledgebase and its supplement TrEMBL in 2003. Nucleic acids research 31, 365–370 (2003).

4. UniProt: the universal protein knowledgebase in 2023. Nucleic Acids Research 51, D523–D531 (2023).

5. M. Ashburner et al., Gene ontology: tool for the unification of biology. Nature genetics 25, 25–29 (2000).

6. S. F. Altschul, W. Gish, W. Miller, E. W. Myers, D. J. Lipman, Basic local alignment search tool. Journal of molecular biology 215, 403–410 (1990).

7. A. Bairoch, PROSITE: a dictionary of sites and patterns in proteins. Nucleic acids research 19, 2241 (1991).

8. A. Krogh, M. Brown, I. S. Mian, K. Sjölander, D. Haussler, Hidden Markov models in computational biology: Applications to protein modeling. Journal of molecular biology 235, 1501–1531 (1994).

9. S. R. Eddy, Profile hidden Markov models. *Bioinformatics (Oxford*, England) 14, 755–763 (1998).

10. M. Blum et al., The InterPro protein families and domains database: 20 years on. Nucleic acids research 49, D344–D354 (2021).

11. M. Kulmanov, R. Hoehndorf, DeepGOPlus: improved protein function prediction from sequence. Bioinformatics 36, 422–429 (2020).

12. M. Kulmanov, R. Hoehndorf, DeepGOZero: improving protein function prediction from sequence and zero-shot learning based on ontology axioms. Bioinformatics 38, i238–i245 (2022).

13. R. You, S. Yao, H. Mamitsuka, S. Zhu, DeepGraphGO: graph neural network for large-scale, multispecies protein function prediction. Bioinformatics 37, i262–i271 (2021).

14. W.-L. Chiang et al., in Proceedings of the 25th ACM SIGKDD international conference on knowledge discovery & data mining. (2019), pp. 257–266.

15. I. M. Nooren, J. M. Thornton, Diversity of protein–protein interactions. The EMBO journal 22, 3486–3492 (2003).

16. D. Szklarczyk et al., STRING v11: protein–protein association networks with increased coverage, supporting functional discovery in genome-wide experimental datasets. Nucleic acids research 47, D607–D613 (2019).

17. A. Krizhevsky, I. Sutskever, G. E. Hinton, Imagenet classification with deep convolutional neural networks. Advances in neural information processing systems 25, (2012).

18. T. Sanderson, M. L. Bileschi, D. Belanger, L. J. Colwell, ProteInfer, deep neural networks for protein functional inference. Elife 12, e80942 (2023).

19. A. Vaswani et al., Attention is all you need. Advances in neural information processing systems 30, (2017).

20. H. Zhang, H. Song, S. Li, M. Zhou, D. Song, A survey of controllable text generation using transformer-based pre-trained language models. ACM Computing Surveys 56, 1–37 (2023).

21. Y. Liu et al., A survey of visual transformers. IEEE Transactions on Neural Networks and Learning Systems, (2023).

22. N. Brandes, D. Ofer, Y. Peleg, N. Rappoport, M. Linial, ProteinBERT: a universal deep-learning model of protein sequence and function. Bioinformatics 38, 2102–2110 (2022).

23. A. Kabir, A. Shehu, GOProFormer: A Multi-Modal Transformer Method for Gene Ontology Protein Function Prediction. Biomolecules 12, 1709 (2022).

24. Z. Lin et al., Language models of protein sequences at the scale of evolution enable accurate structure prediction. BioRxiv 2022, 500902 (2022).

25. J. N. Clifford et al., BepiPred-3.0: Improved B-cell epitope prediction using protein language models. Protein Science 31, e4497 (2022).

26. J. Qiu et al., Large ai models in health informatics: Applications, challenges, and the future. IEEE Journal of Biomedical and Health Informatics, (2023).

27. G. Ahdritz et al., OpenFold: Retraining AlphaFold2 yields new insights into its learning mechanisms and capacity for generalization. bioRxiv, 2022.2011. 2020.517210 (2022).

28. R. Taylor, et al., Galactica: A large language model for science. *arXiv preprint arXiv:2211.09085*, (2022).

29. Y. Fang, et al., Mol-instructions: A large-scale biomolecular instruction dataset for large language models. *arXiv preprint arXiv:2306.08018*, (2023).

30. A. I. Basbaum, D. M. Bautista, G. Scherrer, D. Julius, Cellular and molecular mechanisms of pain. Cell 139, 267–284 (2009).

31. L. McInnes, J. Healy, J. Melville, Umap: Uniform manifold approximation and projection for dimension reduction. *arXiv preprint arXiv:1802.03426*, (2018).

32. J. P. Nataro, J. B. Kaper, Diarrheagenic escherichia coli. Clinical microbiology reviews 11, 142–201 (1998).

33. Á. T. Kovács, Bacillus subtilis. Trends in microbiology 27, 724–725 (2019).

34. J. S. Weitz et al., Phage–bacteria infection networks. Trends in microbiology 21, 82–91 (2013).

35. G. Kozlov et al., Solution structure of the catalytic domain of RICH protein from goldfish. The FEBS Journal 274, 1600–1609 (2007).

36. B. S. Remus, A. Jacewicz, S. Shuman, Structure and mechanism of E. coli RNA 2 ′, 3 ′ -cyclic phosphodiesterase. Rna 20, 1697–1705 (2014).

37. A. Hofmann, M. Grella, I. Botos, W. Filipowicz, A. Wlodawer, Crystal Structures of the Semireduced and Inhibitor-bound Forms of Cyclic Nucleotide Phosphodiesterase from Arabidopsis thaliana* 210. Journal of Biological Chemistry 277, 1419–1425 (2002).

38. A. Q. Jiang, et al., Mistral 7B. arXiv preprint arXiv:2310.06825, (2023).

39. J. Devlin, M.-W. Chang, K. Lee, K. Toutanova, Bert: Pre-training of deep bidirectional transformers for language understanding. arXiv preprint arXiv:1810.04805, (2018).

40. J. Li et al., Align before fuse: Vision and language representation learning with momentum distillation. Advances in neural information processing systems 34, 9694–9705 (2021).

41. B. Buchfink, C. Xie, D. H. Huson, Fast and sensitive protein alignment using DIAMOND. Nature methods 12, 59–60 (2015).

42. C.-Y. Lin, in Text summarization branches out. (2004), pp. 74–81.

43. M. Kulmanov et al., Deepgo-se: Protein function prediction as approximate semantic entailment. bioRxiv, 2023.2009.2026.559473 (2023).

44. P. Radivojac et al., A large-scale evaluation of computational protein function prediction. Nature methods 10, 221–227 (2013).

45. M. Kulmanov, W. Liu-Wei, Y. Yan, R. Hoehndorf, El embeddings: Geometric construction of models for the description logic el++. arXiv preprint arXiv:1902.10499, (2019).

46. V. Gligorijević et al., Structure-based protein function prediction using graph convolutional networks. Nature communications 12, 3168 (2021).

47. X. Shi et al., Convolutional LSTM network: A machine learning approach for precipitation nowcasting. Advances in neural information processing systems 28, (2015).

48. S. K. Burley et al., Protein Data Bank (PDB): the single global macromolecular structure archive. Protein crystallography: methods and protocols, 627–641 (2017).

49. T. N. Kipf, M. Welling, Semi-supervised classification with graph convolutional networks. arXiv preprint arXiv:1609.02907, (2016).

50. L. Zheng et al., AnnoPRO: a strategy for protein function annotation based on multi-scale protein representation and a hybrid deep learning of dual-path encoding. Genome Biology 25, 41 (2024).

51. S. Unsal et al., Learning functional properties of proteins with language models. Nature Machine Intelligence 4, 227–245 (2022).

52. Y. Jiang et al., An expanded evaluation of protein function prediction methods shows an improvement in accuracy. Genome biology 17, 1–19 (2016).

53. M. Kulmanov, M. A. Khan, R. Hoehndorf, DeepGO: predicting protein functions from sequence and interactions using a deep ontology-aware classifier. Bioinformatics 34, 660–668 (2018).

54. W. T. Clark, P. Radivojac, Information-theoretic evaluation of predicted ontological annotations. Bioinformatics 29, i53–i61 (2013).

